# Comparative phylogenetic analyses of limestone and non-limestone Begonias in Peninsular Malaysia

**DOI:** 10.64898/2026.09.10.750513

**Authors:** Ruth Kiew, Fathmath Shaman Fareed, Lian Chee Foong, Sheh May Tam

**Affiliations:** Forest Research Institute Malaysia, 52109 Kepong, Selangor, Malaysia; School of Biosciences, Taylor’s University, Jalan Taylors, 47500 Subang Jaya, Selangor Darul Ehsan, Malaysia; State Key Laboratory of Systems Medicine for Cancer, Department of Urology, Ren Ji Hospital, Shanghai Cancer Institute, Shanghai Jiao Tong University School of Medicine, Shanghai 200127, China (Present address); ex-School of Biosciences, Taylor’s University, Jalan Taylors, 47500 Subang Jaya, Selangor Darul Ehsan, Malaysia

**Keywords:** *Begonia*, limestone, biodiversity, phylogenetics, Malaysia

## Abstract

Karsts of Southeast Asia cover ∼400,000 km^2^, with tropical limestone karsts possessing highly biodiverse flora in many, varied microhabitats that are unique from lowland- and hill forests. *Begonia* L. is a pantropical mega-genus with approximately 2000 described species to date, ideal for studying evolutionary patterns and factors driving tropical population- diversity and speciation. Begonias grow on multiple substrates including limestone and granite, sandstone, quartzite, on steep earth slopes and near streams in primary forests. Niche partitioning is commonly observed, resulting in many single-site endemic *Begonia* species especially on limestone karsts. Molecular phylogenetic analyses were performed on combined chloroplast *ndhF-rpl32* and nuclear internal transcribed spacer *ITS* sequences that included 30 selected *Begonia* species representing all sections from Peninsular Malaysia (Pen. Msia; ten limestone-, thirteen forest- and seven granite species). Results of Bayesian inference and molecular dating showed evolution of *Begonia* from continental Asia into Pen. Msia during the mid-late Miocene period of ∼ 10.65 mya (9.7 – 12.12 mya), with at least two independent dispersal events observed by the early splitting of the ancestral lineage into two major clades i.e. clades A- and B at 9.85 mya (HPD 9.7–10.36 mya) and 8.85 mya (HPD 7.7–11.23 mya) respectively; followed by multiple speciation events colonizing limestone- as well as non-limestone (forest, granite) habitats. *Begonia* speciation into limestone habitats occurred three times, with climate indicated as more important factor than substrate in diversification - clade A contained the “Indian/Continental Asia” (monsoon climate) species, i.e. Sect. Platycentrum and Parvibegonia, resolved slightly older than clade B where Sect. Petermannia, Jackia and Ridleyella formed a “Sunda Shelf” (equatorial climate) centred group. Strong population structuring by locality, common in Begonias, was also observed (e.g. *B. kingiana, B. foxworthyi, B. nurii, B. sinuata*), regardless of distribution. However, for widespread species (*B. kingiana, B. sinuata)* “behavioural (physiological) inertia” on niche evolution, and/or genome size variation and dynamics (genome evolution) may potentially explain species cohesion. Phylogenetic relationships are mostly concordant with current sectional classifications excepting Sect. Parvibegonia, Sect. Diploclinium and two paraphyletic species complexes. Conservation of Begonias and biodiverse limestone karsts in Malaysia remains concerning due to ongoing threats (large-scale quarrying/mining, human encroachment), and most are still without any legal protection.

## Introduction

Asia contains ∼8.35 million km^2^ of karstic habitat, with Southeast Asia covering an area of around 400,000 km^2^ with geological ages ranging from the Cambrian to the Quaternary, being home to rich flora & fauna that exhibit high endemism (Clements et al., 2006; Day and Urich, 2000; Grismer et al., 2021). Limestone karsts are sedimentary rocks consisting of calcium carbonate, mostly formed millions of years ago by calcium-secreting organisms such as corals and lifted above sea level due to tectonic movements. Of high scientific interest, limestone flora (and fauna) is unique from other lowland forest types, and extremely diverse in comparison with the small area it occupies (Chin, 1977; Saw, 2010). Karst topography usually consists of fragmented hills, caves, and towers forming “island-like” habitats across broad geographic areas, and together with their fractured and eroded surfaces, provide many and varied microhabitats that might have contributed to the extraordinarily high degrees of range-restricted endemism (Grismer et al., 2021; R. Kiew et al., 2023; Kiew and Rahman, 2021; Tan et al., 2024).

Karsts in Southeast Asia (SE Asia) are threatened by modern destructive practices, mainly large-scale operations involving quarrying/mining for limestone and basement minerals (Kiew, 1995)(CPD), (Kiew, 1997)(CAP), (Clements et al., 2006; Davison, 2001), encroachment and fires resulting from human activities (R. Kiew et al., 2023). Destruction by quarrying is permanent and irreversible, now regarded as the primary threat to the survival of karst-associated species; only about 13% of SE Asia’s karst area has been nominally protected (Day and Urich, 2000). In Peninsular Malaysia (Pen. Msia), there are about 445 limestone outcrops (Liew et al., 2016; Price, 2014) that cover approx. 0.2% of the Peninsula’s land area. Most karsts are small with a basal area of about one km^2^ and ascend to less than 700 m above the surrounding area but are nevertheless outstandingly biodiverse - harboring about 14% of the Peninsular’s flora (at least 1216 vascular species) (Chin, 1977) of which 21.4% are endemic and 20.8% are restricted to living on limestone.

The renowned botanist E.J.H. Corner first drew attention to the study of mega-genera of flowering plants highlighting how highly speciose genera may contribute to deeper understandings on plant evolutionary biology in tropical forests (Corner, 1961; Frodin, 2004; Moonlight et al., 2018). Interestingly, Begonias in Pen. Msia are highly endemic (89%) and grow naturally on both non-limestone (granite, sandstone, quartzite) and limestone habitats, as well as steep earth slopes in primary rainforests near streams (Chua et al., 2009; Kiew, 2005; Phutthai and Hughes, 2024). *Begonia* L. has emerged as a *bona fide* pantropical mega-genus, currently ranked the sixth-largest flowering plant genus with 70 taxonomic sections and over 2000 described species of mostly perennial herbs or soft wooded shrubs (Frodin, 2004; Goodall-Copestake et al., 2010; Moonlight et al., 2018; Soulivanh et al., 2020). Previous studies had reported on the origin- and migration of Begonias out of Africa (Goodall-Copestake et al., 2010; Plana et al., 2004) during the Oligocene via India (Rajbhandary et al., 2011) and diversifications in Asia in the Miocene period 18-15 Mya (Chung et al., 2014; Hughes et al., 2015; Thomas et al., 2012). Dispersals of Asian Begonias between continental Asia and Malesia had been from west to east, studies dated diversification in the Malesian region and the Philippines to 13.0 Mya (Thomas et al., 2012) and 16.2 Mya (Hughes et al., 2015) respectively; the main driving forces of divergence identified as microallopatry (topographical heterogeneity) for Begonias in Malesia (R. Kiew et al., 2023; Kiew and Rahman, 2021; Tan et al., 2024; Thomas et al., 2012) and limited pollen- & seed dispersal (Chan et al., 2019; Kiew, 2005).

Peninsular Malaysia recorded 53 indigenous *Begonia* taxa currently classified into seven sections *sensu* Moonlight *et al*., 2018 (also Hughes, Girmansyah and Jack, 2011), *i.e.* sect. Platycentrum (including Sect. Sphenanthera, 24 species growing mainly on granite rocks in small streams in rainforests), Jackia (11 species mostly limestone), Parvibegonia (9 species, pre-adapted to dry seasons, grows on granite and limestone), Petermannia (4 species), Ridleyella (2 species), Diploclinium (2 species) and Bracteibegonia (1 species) (Kiew, 2005). On limestone in particular, six species are restricted to moist base, semi-deeply shaded vertical cliff faces (*B. ignorata, B. jiewhoei, B. jayaensis, B. kingiana, B. nurii* and *B. tigrina*), whereas four species, namely *B. phoeniogramma, B. integrifolia, B. variabilis,* and *B. foxworthyi* grow on both limestone as well as bare earth slopes/banks among granite rocks or sandstone substrates (Chua et al., 2009; Kiew, 2005; Kiew and Sang, 2007; Phutthai et al., 2009). Pen. Msia’s Begonias exhibit three habit types exemplified by species in (i) Sect. Parvibegonia - with tuber, stem succulent, low stature (adapted to drought periods); (ii) Sect. Jackia - rhizomatous, stem short with crowded internodes often producing a rosette habit, good for vertical rocks or earth banks, low stature; and (iii) Sect. Petermannia - rhizomatous, stem and internodes long, base of stem woody, “cane-like” tall 1-3 m. Only one species, *B. sinuata*, is strictly annual, although several species in Sect. Parvibegonia are short-lived, dying down in dry periods. Majority of Begonias, including the rhizomatous species are perennial in equatorial climate. Four types of fruits are distinguished (i) fleshy fruit, such as seen in the most widespread species *B. longifolia*, which are presumably eaten by animals; (ii) dry capsules whereby seeds are released through slits between the locules; (iii) splash-cup - fruit where two shorter wings face upwards to form a cuplike structure while the much larger fibrous wing hangs down like a keel allowing seeds to be bounced out by the ballistic force of raindrops from the tree canopy and (iv) capsule with equal-sized wings or with one larger wing, with seeds probably released when the plant/capsule is shaken (Kiew, 2005). Seeds of Begonias are numerous and very small ‘dust seeds’ (0.2-0.5 mm long), making them very unlikely to be dispersed far from the mother plant since they grow in windless habitats e.g. shaded habitats in forest. Seeds are distinctly sculptured with rough surface, perhaps aiding in clinging on to rough rock/cliff surfaces. Being produced from such small seeds, the seedlings are minute, surviving on vertical surfaces of rocks or steep earth banks to avoid coverage by fallen leaves or debris.

Niche partitioning is observed, with many local limestone hills occupied by at least one *Begonia* species, often growing with a second species in the same locality, *e.g*., at Batu Caves, Selangor *B. kingiana* grows on higher vertical rock cliffs while *B. phoeniogramma* is found on soil in deep shade near the base of the hill (Kiew, 2005, 1998). A total of 45.3% (24 of 53 spp.) of Begonias are classified as Critically Endangered, Endangered, or Vulnerable according to International Union for Conservation of Nature (IUCN) conservation categories (IUCN 2019). Only four species have relatively widespread distributions – *B. longifolia* is in India, China, Vietnam, Thailand, Pen. Msia, and Indonesia (Sumatra and Java); *B. martabanica* in Myanmar, Thailand and Pen. Msia (Perak); *B. sinuata* in Vietnam, Thailand and throughout Pen. Msia, whilst *B. integrifolia*’s distribution ranges from India, Thailand, Myanmar to Pen. Msia (Kiew, 2005). Most widespread amongst limestone species is the endemic *B. kingiana*, in comparison to other species such as *B. jayaensis, B. jiewhoei, B. nurii* and *B. tigrina* that are only found in the single state of Kelantan (Kiew, 2005). Most Begonias are known to have very narrow distribution ranges (Hughes and Hollingsworth, 2008; Kiew, 2005; Kiew and Sang, 2015; Tebbitt et al., 2006), hence records of single-site endemic species are fairly common, especially on limestone karsts (Dewitte et al., 2011; Kiew, 2019, 2001; Peng et al., 2008; Tan et al., 2024). For *Begonia* sect. Coelocentrum and allied limestone species found on Sino-Vietnamese karsts, Chung *et al*. (2014) reported a single evolutionary origin of the adaptation to limestone substrates by rhizomatous *Begonia* spp., followed by subsequent species radiation characterized by a strong tendency for niche conservatism. This is in concordance with proposed allopatric speciation linked to palaeogeological events and ancestral periods of climatic fluctuations as drivers of speciation, leading to the diversification of Begonias in SE Asia (Thomas *et al*., 2011, 2012; Chung *et al*., 2014 and references therein).

Karsts remain severely understudied (Dennis and Aldhous, 2004; Sang and Kiew, 2014; Tan et al., 2024; Vermeulen and Whitten, 1999), and on the highly fragmented Sunda Shelf, karsts have formed “islands within islands” with groups of small and isolated karsts observed to contain high levels of endemic flora (Kiew et al., 2023; Kiew and Sang, 2015; Tan et al., 2024). With more than 550 *Begonia* species distributed in SE Asia (Hughes and Hollingsworth, 2008; Thomas et al., 2011), much remains to be studied to gain deeper understanding into the evolution/diversification of the megadiverse *Begonia* in this region. In this comparative molecular phylogenetic study of selected 30 limestone- and non-limestone (forests, granite) species of *Begonia* in Pen. Msia, main findings showed evolution of Begonia from continental Asia into Peninsular Malaysia during the mid-late Miocene period of ∼ 10.65 mya, with at least two independent dispersal events observed, followed by multiple speciation events colonizing limestone- as well as forest-, granite habitats. *Begonia* speciation into limestone habitats occurred three times, with climate being a more important factor than substrate in Begonia diversification, with early, two major clades containing species of “Indian/Continental Asia” (monsoon climate), i.e. Sect. Platycentrum and Parvibegonia, and “Sunda Shelf” (equatorial climate) i.e. Sect. Petermannia, Jackia and Ridleyella affiliations respectively. Common for Begonias, strong population differentiation was detected, as well as surprisingly, species cohesion for mid-aged, widespread species (*B. kingiana, B. sinuata*) perhaps caused by “physiological inertia” on niche evolution, and/or genome dynamics. Phylogenetic results are concordant with previous studies for Begonias in Southeast Asia (Malesia), contributing to the growing knowledge bank which will be useful for Begonias and limestone biodiversity conservation and practice.

## Material and Methods

### Material

Representative taxa from all seven *Begonia* sections from Pen. Msia *sensu* Moonlight *et al*., (2018), totalling 30 species (177 samples) that included all ten limestone species, thirteen non-limestone (forest-) and seven granite species were mostly sampled from the living collections of the Forest Research Institute of Malaysia (FRIM). Additional samples were collected via fieldwork at Batu Caves and the Gabai Waterfalls (Selangor), Sekayu Recreational Forest (Terengganu), Hutan Simpan Pasoh and Lata Tingkat (Negeri Sembilan) and Lanchang Forest Recreational Park (Pahang) in Pen. Msia (December 2018 to January 2020) (Supplementary file 1). For each sample, a small amount of young leaf material was cleaned, cut into smaller pieces and dried using silica gel. Voucher specimens were made and deposited at the national herbarium, Forest Research Institute Malaysia (KEP).

### DNA Extraction, DNA Quantification, PCR amplification and DNA Sequencing DNA Extraction and DNA Quantification

Total DNA was extracted from silica dried young leaf samples using the NucleoSpin® Plant II kit following the manufacturer’s protocol (Macherey Nagel, Germany). Modifications to the protocol were made for certain samples that included increasing the volume of lysis buffer added (up to 2.5x) and reducing final DNA elution volume to 40 μL of DNAse/RNase free water (Fisher Scientific, U.S.A.). All DNA samples were stored at -20°C until further use. Optical Density (OD) readings at wavelengths 230nm, 260nm, and 280nm were recorded for each sample using a UV-Visible spectrophotometer (Beckman Coulter, U.S.A). Samples that did not meet OD_260/230_ ratio of 1.8 to 2.2, and the OD_260/280_ ratio of 1.8 to 2.0 (indicating low purity) were subjected to DNA re-extraction.

### Polymerase Chain Reaction (PCR) Amplification

Two DNA marker regions namely the chloroplast DNA ndhF-rpl32 region (Thomas et al., 2012, 2011) and the nuclear ribosomal DNA internal transcribed spacer (ITS) region (Chung et al., 2014; Tebbitt et al., 2006) were used in this study, based on their proven utility from previous *Begonia* phylogenetic studies. For the ndhF-rpl32 marker, PCR amplification was performed in 25 μl total volume: 12.5 μl GoTaq 2x Promega DNA Master Mix (Promega Corporation, U.S.A.), 1.0 μl of each forward and reverse primer (10 μM) (Integrated DNA Technologies, U.S.A) (Supplementary file 2), 1.0 μl BSA (Bovine Serum Albumin) (New England BioLabs Inc., U.S.A.), 2.0 μl MgCl_2_ (Promega, U.S.A.), 7.5 μl Nuclease free water (Promega Corporation, U.S.A.) and 2.0 μl of total DNA template (∼ 100 ng). The addition of PCR additives such as 1.0 μl BSA and/or 2.0 μl MgCl_2_ were required to overcome PCR inhibitors, enhance PCR amplification success and yield. For the ITS marker, amplification reactions were similarly performed in a total volume of 25 μl with the addition of 0.1 μl extra Taq, (GoTaq Flexi DNA polymerase), 1.0 μl of DMSO (Dimethyl Sulfoxide) (Sigma Aldrich, Malaysia), 1.0 μl Betaine (Sigma Aldrich, Malaysia), 1.0 μl BSA (New England BioLabs Inc., U.S.A.), 2.0 μl MgCl_2_ (Promega, U.S.A.), 5.4 μl Nuclease free water (Promega Corporation, U.S.A.) and 2.0 μl of total DNA template (∼ 100 ng). The addition of Betaine and an extra 0.1 μl of Taq polymerase, along with DMSO and MgCl_2_ were effective in ensuring PCR success. PCR reactions were performed for 40 cycles generally following the protocols by Thomas et al., (2011) for the ndhF-rpl32 region and Chung et al., (2014) for the ITS region, with minor adjustments (Supplementary file 3).

### Visualization of PCR amplicons and DNA sequencing

Agarose gel 1.0% (w/v) was prepared by melting 0.3g of agarose powder (Fisher Scientific, U.S.A.) in 30.0 mL of 1X Tris-Borate-EDTA (TBE) buffer (1st Base, Singapore), with the addition of 3 μL of SYBR Safe DNA Gel Stain (Invitrogen, U.S.A.). PCR products (5 μL) were loaded on to the 1.0% agarose gel, and electrophoresis was performed at 80V for ∼30 mins. Agarose gel visualization was performed using the UV Imager (Vilber Lourmat Deutschland GmbH, Germany). The expected band size for both ndhF-rpl32 and ITS amplicons was approximately 800 bp, and one clear band taken to indicate successful amplification. Band sizes were compared using Gene Ruler 1 kb Plus DNA Ladder (Fisher Scientific, U.S.A.).

PCR products of samples that showed one clear expected band size were purified using Wizard® SV Gel and PCR Clean-Up System kit (Promega Corporation, U.S.A.) following the manufacturer’s instructions. Purified samples were then sent for DNA sequencing using the Single Pass DNA sequencing method using forward and reverse primers of both *ndhF-rpl32* and *ITS* respectively, performed by First Base Laboratories Sdn. Bhd., Selangor, Malaysia.

### Phylogenetic Analyses Multiple Sequence Alignment

Consensus sequences for each sample were generated using the program BioEdit (Hall, 1999) and then subjected to Blastn (https://blast.ncbi.nlm.nih.gov) against the National Centre of Biotechnology Information (NCBI) nucleotide database for verification (Begonia species; correct DNA marker region). Multiple sequence alignment of generated *Begonia* consensus sequences together with two African outgroup sequences, i.e. *B. dregei* (AY429336.1) and *B. sutherlandii* (AF485215.1) (Chung et al., 2014) were performed on BioEdit (Hall, 1999) using the ClustalW Multiple Alignment feature. Ambiguous data at the beginning and end of the dataset were trimmed, but the gaps resulting from insertions and deletions (indels) in the aligned data matrix were retained. The combined aligned matrix consisted of 156 taxa with 2161 aligned characters (Table 1). Partition homogeneity test (incongruence length difference test, ILD) was performed using PAUP* (Swofford, 2002) with 1000 homogeneity replicates of heuristic searches with ten random sequence additions and tree-bisection-reconnection (TBR) branch swapping, following the parameters used in the study by Chung et al., (2014). All characters were unordered and had equal weight, and gaps were treated as missing data.

**Table 1:**
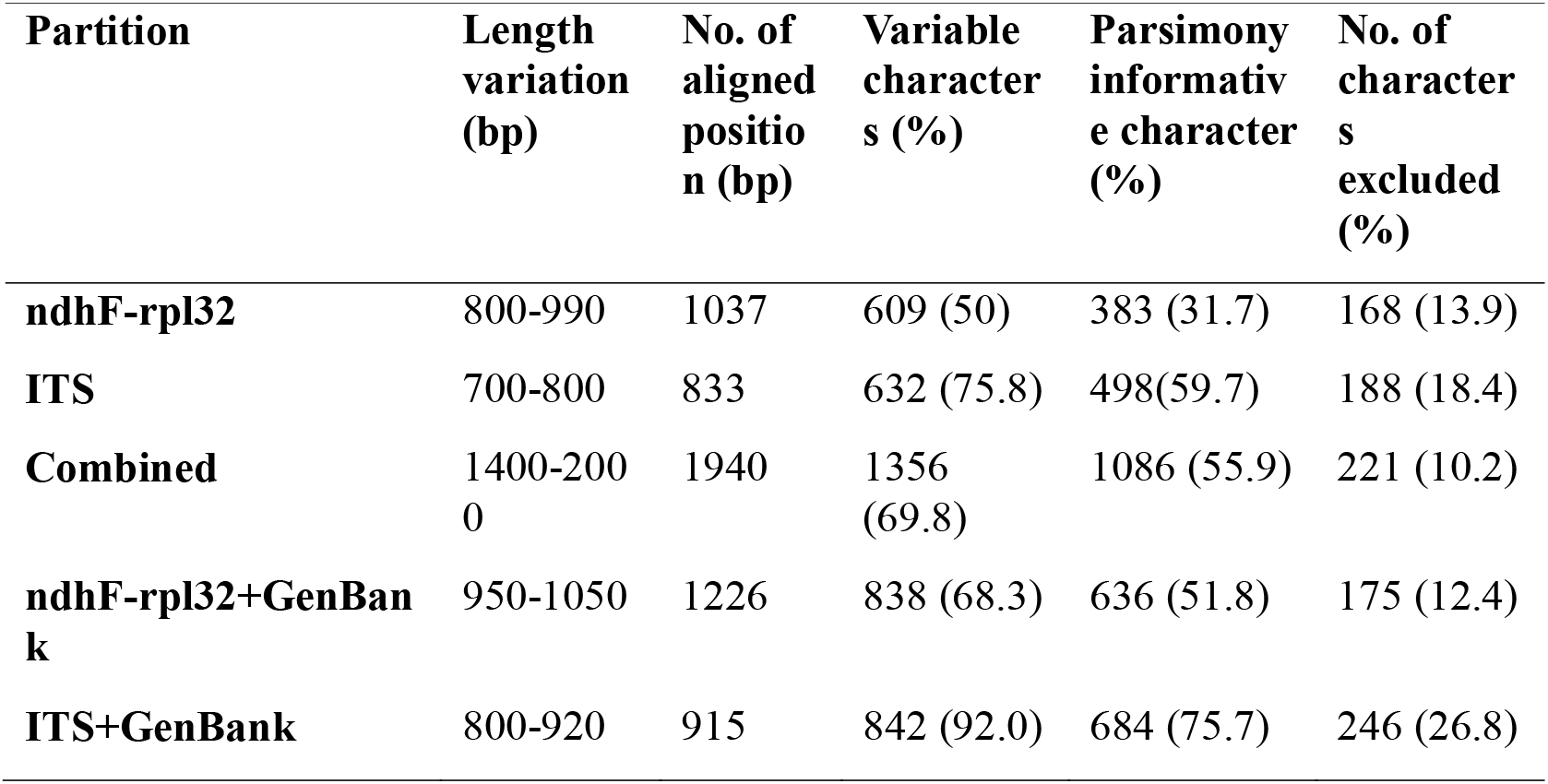
Sequence information for chloroplast ndhF-rpl32 (161 samples), nuclear ITS (156 samples) and the two combined DNA regions for Begonia samples used in this study. Characters were unordered, had equal weight and gaps were treated as missing data

### Bayesian Inference Analysis

Nucleotide substitution models were analysed using jModelTest2 on XSEDE via the CIPRES portal (Posada, 2008) according to Akaike Information Criterion (AIC/AICc - correction for small sample sizes) for the ndhF-rpl32 dataset was TVM+G (transversional model) with among-site variation distribution. However, this model was not implemented by MrBayes, the GTR+G (General Time Reversible) model for nucleotide substitution + allowing for rate variation for gamma distribution among sites was applied, following Thomas et al., (2011). Models selected by AIC/AICc for ITS region were TVM+I+G and K80+G (Kimura’s two-parameter) respectively, hence the K80+G was implemented for Bayesian analysis. For the combined dataset, both AIC/AICc selected the TIM2+I+G (transitional model), but again the GTR+G model was implemented.

Bayesian phylogenetic reconstructions were performed through the CIPRES portal using MrBayes v3.2.6 on XSEDE (Darriba et al., 2012; Guindon and Gascuel, 2003). Each search used three incrementally heated and one MCMC (Markov Chain Monte Carlo) with a temperature setting of 0.2 and was run for 1 x 10^6^ generations, number of runs = 2, number of chains = 4, with every 1000 generations sampled and burn-in fraction of 0.25. All the other parameters were kept on the default setting. Resulting phylograms were viewed using the software FigTree (Rambaut, 2007) using outgroup rooting, depicting 50% majority consensus tree with branch number representing Bayesian posterior probability value. A 95% posterior probability (BPP) was considered as well-supported clades or relationships (Thomas et al., 2011).

## Molecular Dating

The software BEAST2 on XSEDE v2.6.3 via CIPRES portal (Bouckaert et al., 2014; Drummond and Rambaut, 2007) was used to perform molecular divergence time estimates upon formatting the aligned matrix using BEAUti with a single partition using GTR+G model. The phylogenetic tree was calibrated using priors estimated by Thomas et al., (2012) for Asian *Begonia* clade at 16.1 and Malesian *Begonia* clade at 13.0 with a standard deviation of 3.265 (95% (HPD) 9.7-22.5) and 2.7 (95% (HPD) 7.7-18.3) respectively, following Chung et al., (2014). The tree prior was set to Yule process and uncorrelated-rates relaxed molecular clock with a lognormal distribution of rates was utilized. BEAST analysis was run for 50 million generations sampling every 1000th generation with a 10% burn-in (Chung et al., 2014; Thomas et al., 2012). The resulting maximum clade credibility chronogram was viewed using the software FigTree (Rambaut, 2007).

## Results

Summary of *Begonia* sequence information for ndhF-rpl32-, ITS- and combined dataset are presented for 161 samples for ndhF-rpl32 region and 156 samples for the ITS region. The final combined dataset consisted of 156 taxa and 2161 aligned positions whereby 1940 were included in the analyses, of which 1356 (69.8%) were variable characters (Table 1). A total of 221 bp (10.2%) of aligned positions were excluded due to poor alignment mainly at the beginning and end of sequences. Interestingly, all samples of *B. wrayi* had a long deletion in the ndhF-rpl32 region which reached 422 bp.

Partition Homogeneity Test found significant difference (P-value=0.001) between the ndhF-rpl32-, ITS datasets and random partitioning indicating heterogeneity among datasets. Individual BI phylograms inferred by the ndhF-rpl32- and combined dataset were highly similar but not with the ITS dataset. Comparisons showed that differences were not within species relationships (clades resolved) but mainly in basal topology after the initial split from the African outgroup species. Explained as soft incongruences, Chung et al., (2014) had in their study observed similar results, with significant ILD test results (P=0.001) where topology and relationships of the ITS- and combined dataset trees were highly congruent but not with the rpL16 phylogram. Following Chung et al., (2014), since the combined dataset had better resolutions for species relationships and mostly higher values of clade support, the combined dataset is used here for further analyses and discussion.

The combined BI tree with posterior clade probabilities (BPP) of Pen. Msia’s *Begonia* species show an early division into two major clades, labelled as major clades- A (BPP=82) and B (BPP=82) with similar, short branch lengths (0.006) comprising of seventeen- and thirteen species respectively (Figure 1). Clade A consists of fourteen non-limestone species and three species that can be found on both limestone and granite habitats. Two subclades A1 (BPP=100) and A2 (BPP=91) are identified (Figure 2) - A1: a group of four species i.e. *B. integrifolia, B. variabilis, B. elisabethae*, and *B. phoeniogramma* (Sect. Parvibegonia, growing in both limestone and granite habitats) is sister to a group of eight forest species (most grow on granite rocks associated with streams), namely *B. rhoephila, B. rheifolia, B. pavonina, B. herveyana, B. venusta, B. abdullahpieei, B. maxwelliana*, *B. decora* and *B. longifolia* (Sect. Platycentrum).

**Figure 1.**
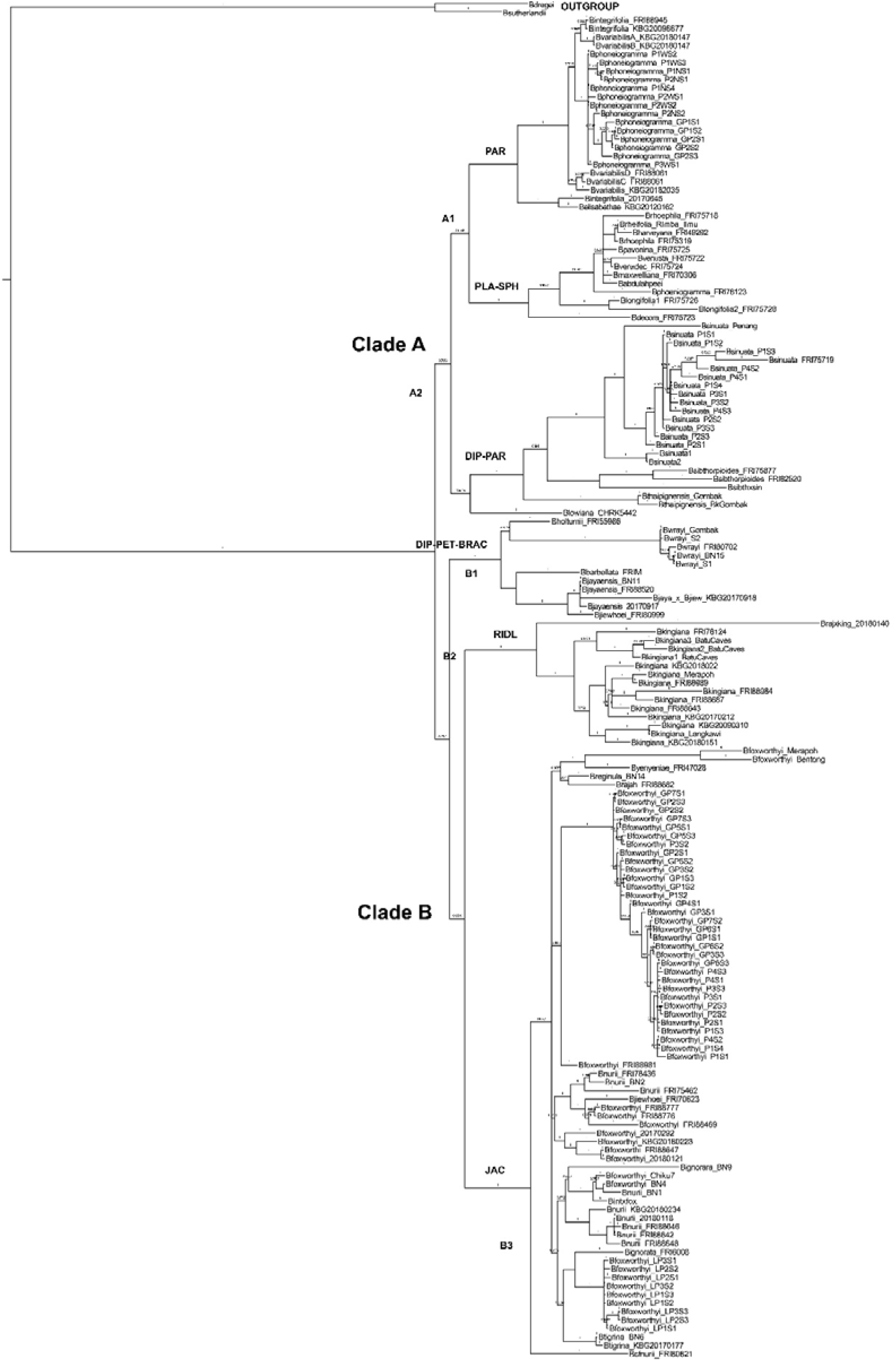
Bayesian inference phylogram of 30 Begonia species (156 taxa) from Peninsular Malaysia based on combined ndhF-rpl32 and ITS sequences. Phylogram shows an early division into two major clades, labelled as Clade A (BPP=82) and clade B (BPP=82). Major clade A (BPP=82) and clade B (BPP=82) comprise of seventeen- and thirteen species respectively. Bayesian posterior probability (BPP) support values are shown above each branch. Labels refer to sections of Begonia: PLA-SPH (Platycentrum-Sphenanthera), PAR (Parvibegonia), DIP (Diploclinium), BRAC (Bracteibegonia), PET (Petermannia), JAC (Jackia), and RIDL (Ridleyella).

**Figure 2.**
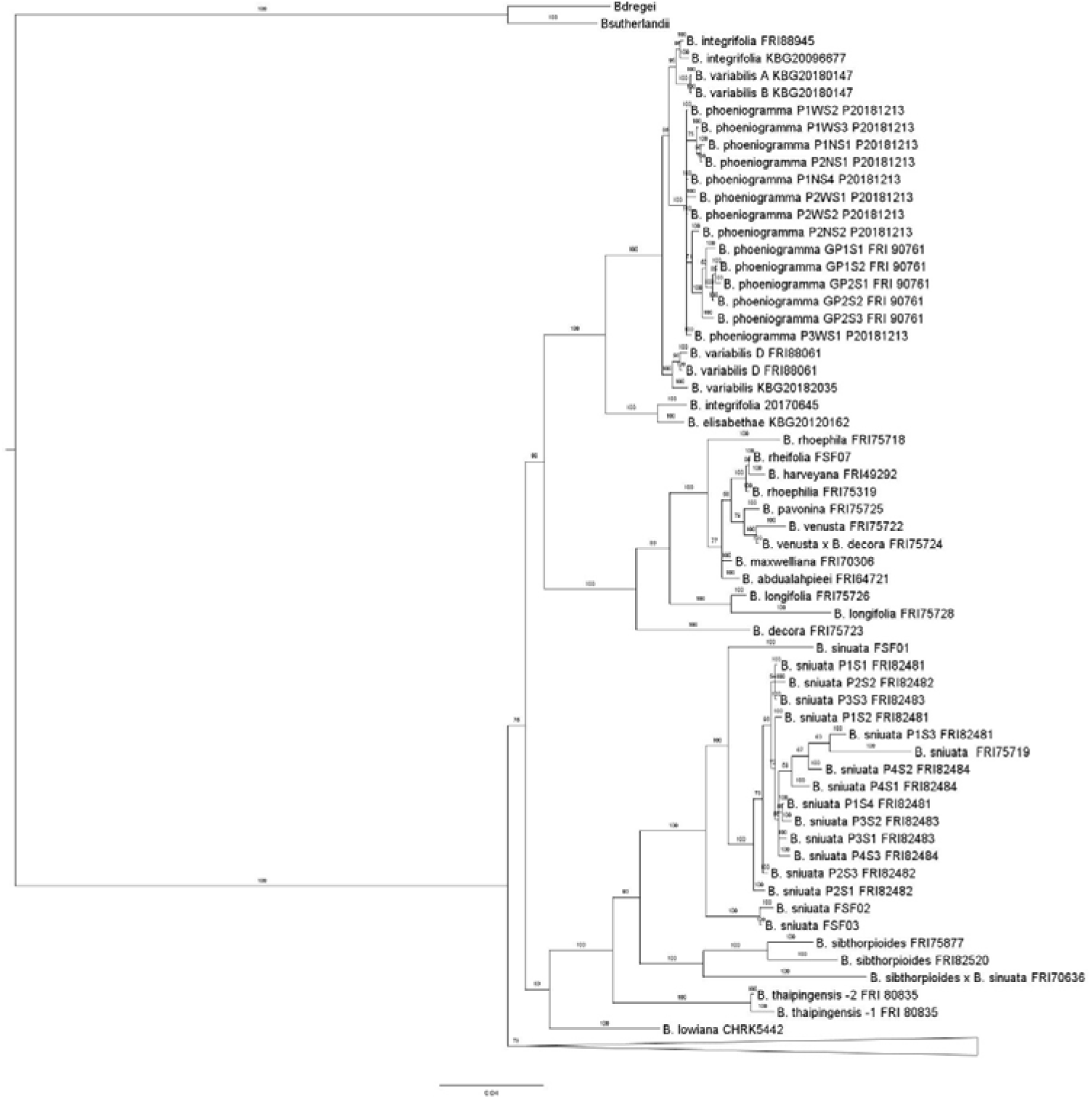
Major Clade A of Bayesian inference phylogram of 30 Begonia species from Peninsular Malaysia based on combined ndhF-rpl32 and ITS sequences. Values above the branches denote branch length. Labels above main branches refer to Sectional classifications – PLA-SPH (Platycentrum-Sphenanthera), PAR (Parvibegonia), and DIP (Diploclinium). Two subclades A1 (BPP=100) and A2 (BPP=91) are further identified.

Interestingly, individuals of *B. phoeniogramma* – labelled as ‘P’ from Batu Caves (limestone) and ‘GP’ from Gabai Waterfalls (granite) have segregated based on locality and not by habitat. In subclade A2, *B. sibthorpioides, B. sinuata* (considered as “*unusual”* species) in Sect. Parvibegonia and their hybrid, resolved as sister species and together formed a sister group to *B*. *thaipingensis*, clearly apart from other “*typical*” Sect. Parvibegonia species (Kiew, 2005). A lone species *B. lowiana* was positioned basal to this group, distant from *B. jayaensis –* both species are currently classified in Sect. Diploclinium. One sample of *B. kingiana* (FRI88984) resolved amongst the *B. sinuata* clade (misplaced), which most probably is due to labelling error (see below).

Clade B contains three well-resolved subclades (labelled as B1, B2 and B3) each with BPP=100, comprising of six limestone restricted species, three non-limestone species and one on both substrates (Figure 3). Subclade B1 – *B. jayaensis* (Sect. Diploclinium) and *B. jiewhoei* (Sect. Petermannia) resolved as sister species, together with *B. barbellata* (Sect. Bracteibegonia), which is sister to the group of *B. holttumii* and *B. wrayi* (Sect. Petermannia). Subclade B2 consists of endemic limestone species *B. kingiana* (Sect. Ridleyella) where individual samples resolved according to regions – Northern states (KBG20180151, FSF16, KBG20090310), East coast (KBG20170212, FRI88643, FRI88687, FRI88984, KBG2018022, FRI88989, FSF17) and West coast (*B. kingiana-*1,2,3-K20181213, FRI78124). This *B. kingiana* subclade also contains one sample of *B. sinuata* (P4S1FRI82484), suggesting mistaken labelling and both implicated samples will be further verified. The artificial hybrid (KBG20180140) resolved basal to all *B. kingiana* samples, with the ndhF-rpl32 phylogram results confirming the parentage of maternal parent being *B. kingiana* and *B. yenyeniae* was the pollen source).

**Figure 3.**
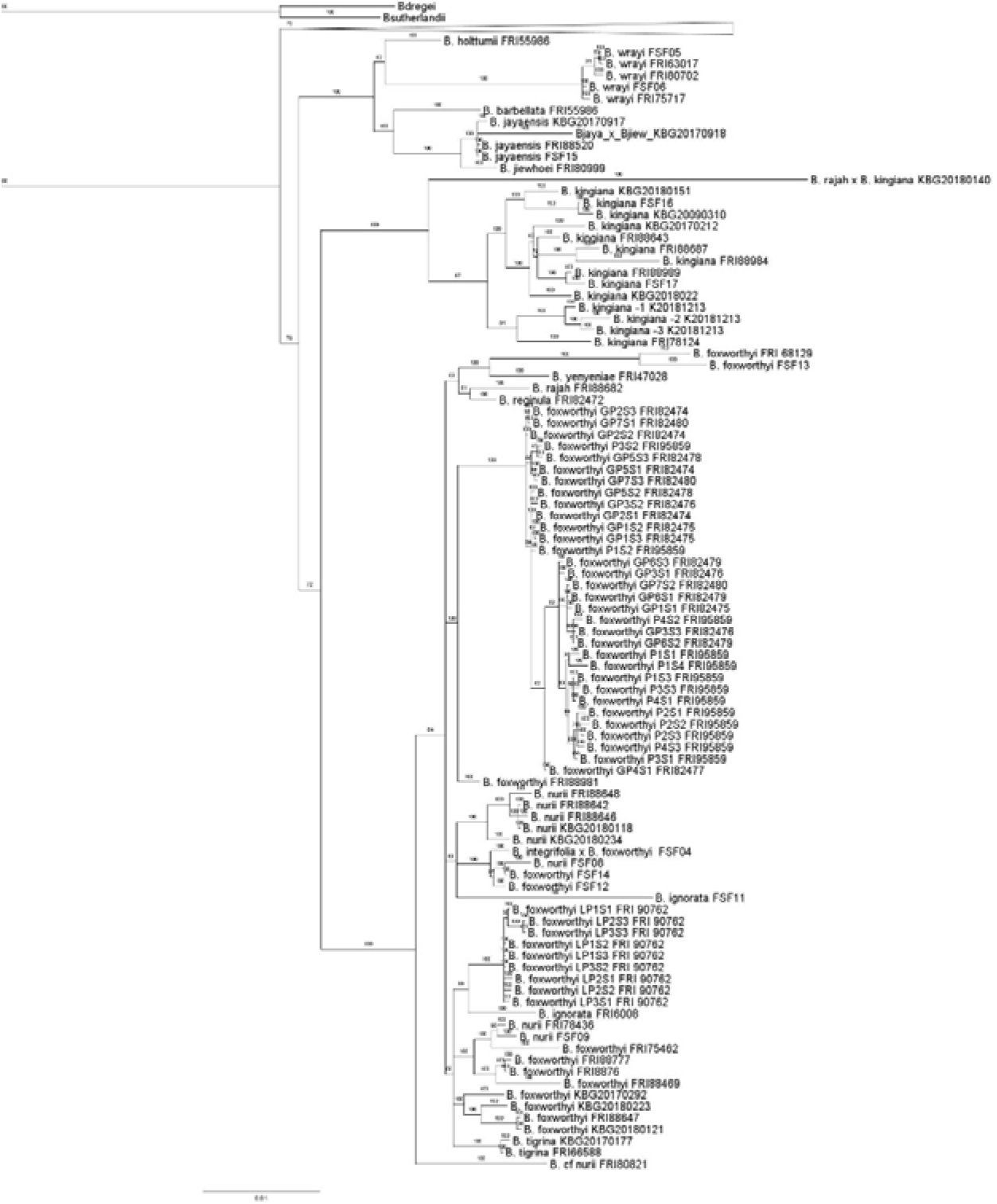
Major Clade B of Bayesian inference phylogram of 30 Begonia species from Peninsular Malaysia based on combined ndhF-rpl32 and ITS sequences. Values above the branches are branch length. Sectional labels above main branches are – DIP (Diploclinium), BRAC (Bracteibegonia), PET (Petermannia), JAC (Jackia), and RIDL (Ridleyella). Clade B contains three well-resolved subclades (B1, B2 and B3) each with BPP=100.

Seven species (Sect. Jackia) formed paraphyletic Subclade B3 with *B. foxworthyi* interspersed among *B. yenyeniae, B. tigrina, B. nurii* and *B. ignorata*. Again, it is shown that *B. foxworthyi* segregated geographically based on locality, with granite samples (‘GP’) from Lata Tingkat, Pasoh, Negeri Sembilan grouping with the limestone samples (‘P’) from Hutan Simpan Pasoh, Negeri Sembilan, removed from other granite samples (‘LP’) from Lanchang, Pahang as well as the limestone *B. foxworthyi* samples from Kelantan.

## Molecular dating results

The maximum clade credibility chronogram and summary of divergence time estimates, and clade supports (PP) generated by the BEAST software are shown in Table 2 and Supplementary file 4. Estimated divergence times of lineages include the entire *Begonia* ingroup during Mid-Late Miocene at 11.33 mya (PP 1; HPD date range 9.7 – 14.26) until the youngest split recorded in Late Pleistocene at 0.06 mya (PP 0.08; HPD date range 0 – 0.2). Clade support for divergence ranged from 1 to 0.04, hence dating results for certain clades/subclades lack confidence (Table 2). Molecular dating results estimated the mean divergent crown age of the ancestral lineage of Begonias (*Begonia* ingroup) of Pen. Msia at 10.65 mya (PP 0.4; HPD 9.7 – 12.12). The earliest split resulting in major Clades A and B showed an estimated crown age of Clade A at 9.85 mya (PP 1; HPD 9.7 - 10.36) and just a slightly younger estimated mean divergent crown age of 8.85 mya (PP 1; HPD 7.7 - 11.23) for major Clade B (Supplementary file 4).

**Table 2:**
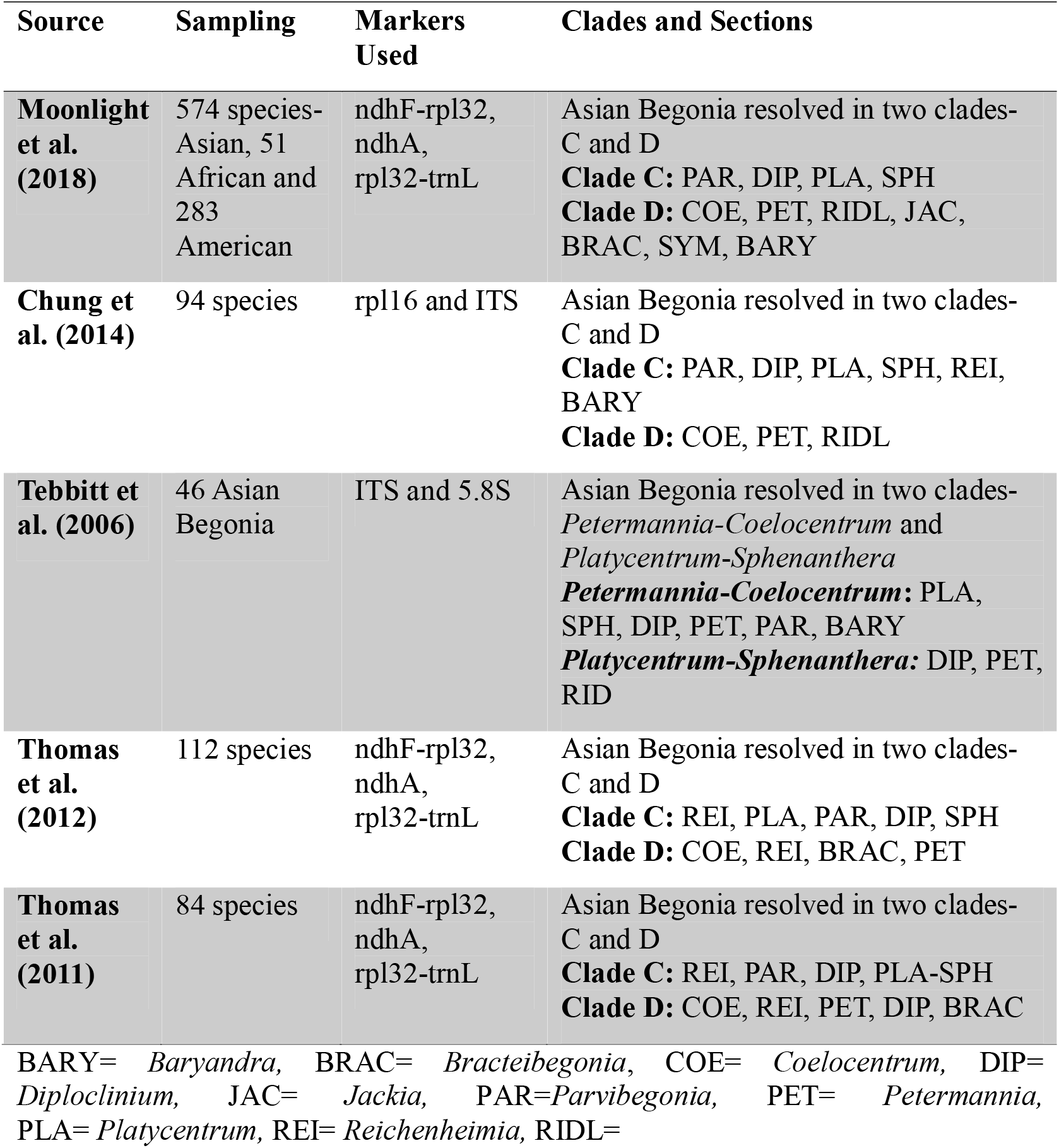
Maximum clade credibility chronogram and summary of divergence time estimates, and clade supports (PP) generated by the BEAST software for Peninsular Malaysian *Begonia* species

In major Clade A, the mean age of origin of the limestone group of species (Sect. Parvibegonia) was estimated at 4.92 mya (PP 1; HPD 2.86 - 7.16) whereas the corresponding subclade of forest-species sister group (Sect. Platycentrum- Sphenanthera) was 5.44 mya (PP 1; HPD 3.31 - 7.7). The split between the non-limestone *B. lowiana* was at 8.46 mya (PP 0.83; HPD 6.47 – 9.78 mya), and emergence of the group of *B. sinuata - B. sibthorpiodes - B. thaipingensis* was dated at 6.63 mya (PP 1; HPD 4.58 - 8.47). Other notable dating estimations include recent emergence of *B. longifolia* at 1.82 mya (PP 1; HPD 0.37 - 3.42); *B. phoeniogramma* limestone population (Batu Caves) at 1.53 mya (PP 1; HPD 0.77 - 2.36) and 0.56 mya (PP 1; HPD 0.21 - 0.97) for the non-limestone population (Gabai Waterfalls) (Figure 2, Supplementary file 4).

In major Clade B, the earliest mean divergent crown age was estimated at 8.04 mya (PP 0.66; HPD 6.01 - 10.59) for subclade B1, within which emergence of limestone species *B. jiewhoei* and *B. jayaensis* was dated at 1.69 mya (PP 1; H HPD DP 0.64 – 3.04). Subclade B2 containing *B. kingiana* had a mean age estimate of 4.62 mya (PP 0.98; HPD 2.93 – 6.43 excluding the artificial hybrid). Subclade B3 which consists of both limestone and non-limestone species complex was dated at 6.09 mya (PP 1; HPD 4.05 – 8.02). Limestone populations of *B. foxworthyi*-*B. nurii* diverged from the granite populations at 3.56 mya (PP 0.44; HPD 2.34 – 4.82). Within the large subgroup of mixed *B. foxworthyi* samples (PP 1; 3.44 mya, HPD 2.01 – 4.95), the granite population was dated at 1.98 mya (PP 1; HPD 1.13 – 2.98) whereas the limestone population was slightly younger at 1.00 mya (PP 0.58; HPD 0.51 – 1.5). Other estimates of mean divergent crown age include *B. ignorata*-*B. foxworthyi* complex of granite population (Lanchang) at 2.33 mya (PP 0.26; HPD 1.0 – 3.71); *B. tigrina*-*B. ignorata*-*B. foxworthyi* at 3.11 mya (PP 0.99; HPD 1.77 – 4.55); *B. foxworthyi* (FRI88469, FRI88776, FRI88777), *B. nurii* (FSF09, FRI78436, FRI75462) at 2.46 mya (PP 0.15; HPD 1.26 – 3.76) and *B. foxworthyi* -*B. yenyeniae*-*B. rajah*-*B. reginula* at 3.59 mya (PP 0.99; HPD 2.27 – 5.04).

## Discussion

### Phylogenetic relationships of Peninsular Malaysian species of *Begonia* inferred from combined *ndhF*-*rpl32* - *ITS* sequences

Phylogenetic relationships of the major clades of limestone and non-limestone *Begonia* species in Pen. Msia inferred from BI of combined dataset (*ndhF-rpl32* - *ITS*) are highly concordant with previous findings such as reported by Tebbitt et al., (2006), Rajbhandary et al., 2011; Thomas et al., (2011, 2012), Chung et al., (2014) and Moonlight et al., (2018) – whereby our sampled Begonias resolved in two major clades (labelled as A and B in this study) that clearly correspond to Asian clades C and D respectively in Thomas et al., (2011) as well as Chung et al., (2014). Results indicate evolution of *Begonia* from continental Asia into Pen. Msia during the mid-late Miocene period of ∼ 10.65 mya (9.7 – 12.12 mya), upon which at least two independent dispersal events were observed in the early splitting of the ancestral lineage into major clades A and B at 9.85 mya (HPD 9.7–10.36 mya) and 8.85 mya (HPD 7.7–11.23 mya) respectively; followed by multiple speciation events in Pen. Msia colonizing limestone- as well as non-limestone (forest, granite) habitats. This key finding aligns with the proposed trend of *Begonia* dispersals between continental Asia and Malesia, in a “west-to-east” manner accompanied by considerable local diversifications (Thomas et al., 2012), plausible because Pen. Msia is the southernmost Southeast Asia continental landmass, which has been continuous for eons (Morley, 2001; Van Welzen et al., 2005). Molecular dating results from our study are concordant to the age estimates findings by Chung et al., (2014) and within the ranges reported by Thomas et al., (2012), albeit being based on different combinations of chloroplast- and nuclear sequence datasets. Thomas et al., (2012) reported divergence age between their major clades C and D at 16.1 mya (9.1–22.5), during the mid-Miocene period with relevant species included specifically in their clade C16 (PP: 1 – dated at 14.3 mya (8.2 – 20.2)) and clade D49 (PP: 1 – 13 mya (7.7 – 18.9)), dominated by continental Asian species (including Indian species) and Malesian/Sunda Shelf species respectively.

Distinct differences in stratigraphy, structure, magmatism, geophysical signatures and geological evolution revealed early formation of three “North–South” belts in Pen. Msia from Late Carboniferous–Early Permian to Late Triassic i.e. the Western-, and Central- & Eastern- belts corresponding to the Sibumasu Terrane, derived from the NW Australian Gondwana margin and the Sukhothai Arc constructed on the margin of the Indochina Block (Metcalfe, 2013). The mountainous terrain of Pen. Msia today are aligned longitudinally from west to east, consisting of eight major ranges i.e. Nakawan, Keda-Singgora, Bintang, Kledang, Main, Benom, Tahan and East Coast ranges (Kasim et al., 2020). Geological barriers may be a plausible explanation for the early splitting and sustained separation during speciation of the ancestral *Begonia* lineage in Pen. Msia. In this study, major clade A contains all the “Asian” species sampled (*B. longifolia, B. integrifolia, B. sinuata*) as well as *B. elisabethae*, one of three species which distribution extends into Pen. Thailand. The other two species, *B. barbellata* and *B. wrayi* resolved in a basal subclade in major clade B. Taken together, phylogenetic results support a trend of dispersal from continental Asia (major clade A, basal clade B) followed by speciation on the Sunda Shelf. In addition, it appears that for Pen. Malaysia, climate was a more important factor than substrate in *Begonia* evolution, as demonstrated by the lithophytic habit being well-represented in both major clades A and B. One clear distinction observed is that clade A contains “Indian/Continental Asia – monsoon climate” species, i.e. Sect. Platycentrum with rhizomatous lithophyte, many non-limestones, speciate especially in mountains (Platycentrum is especially speciose in the Himalayas), and Sect. Parvibegonia, growing on both granite and limestone but morphologically not adapted to lithophytic habitat but instead possesses tubers suited to seasonal “monsoon” climate surviving dry seasons. These species being slightly older than those in major clade B where Sect. Petermannia, Jackia and Ridleyella form a “Sunda Shelf – equatorial climate” centred group with Sect. Petermannia comprises of at least 95% of *Begonia* species found in diverse habitats including limestone in Borneo and the two latter sections are limestone groups.

The majority of species resolved in clade A have non-limestone habit (14 species), with additional three species that grow on both limestone/non-limestone habitats. A mix of limestone restricted (six species), non-limestone (six) and one species that grow on both habitats resolved in clade B. Phylogram topology and molecular dating results (clade A = 9.85 mya compared to clade B = 8.85 mya) indicate the ancestral state being non-limestone (e.g. forest, granite habitats). *Begonia* speciation onto limestone habitats in Pen. Msia occurred mainly three times, as observed once in major clade A – (i) Sect. Parvibegonia with four (of nine) species that are short-lived and die down in dry periods only to regenerate from tubers or seeds at the next rain (*B. integrifolia, B. phoeniogramma, B. variabilis, B. elisabethae*); and twice in major clade B – (ii) *B. jayaensis* (Sect. Diploclinium) and *B. jiewhoei* (Sect. Petermannia) being limestone restricted and (iii) large group of species from Sect. Jackia and Sect. Ridleyella. Sect. Jackia contains a mix of species with rhizome, rosette habit, round leaves, perennial, grow on both rock- as well as limestone habitats (e.g. *B. foxworthyi),* in contrast to the only one species, limestone endemic *B. kingiana* with thick, succulent, peltate leaves, widespread on both east and west of the Main Range belonging to Sect. Ridleyella.

Orogenic events also point to very early formation and presence of limestones karsts in Pen. Msia, such as the well-studied Kinta Valley sequence (Ipoh, Kinta), lying on strike between the Kubang Pasu, Kati and Kenny Hill Formations which represents 700 m of Carboniferous through Lower Permian limestone as well as the Ratburi limestone (southern Thailand) and Chuping Formation in northern Pen. Msia formed during the Artinskian (Hutchison, 2014, 2007). Muhammad and Komoo (2003) reported folding and metamorphosis of limestone and schist during/near the end of Permian, that were then intruded by the Kledang and Main Range granites during Late Triassic, with slow limestone emergence estimated at 0.1 mm per annum (Krähenbuhl, 1991). However, recent studies of Miocene reefs in southern South China Sea suggest possible acceleration and expansion of limestone karst exposure in this region during mid-late Miocene period - coinciding with the estimated colonization period of *Begonia* ancestral lineage into Pen. Msia, due to abrupt onset- and intensification of Asian summer monsoons, namely the East Asian Summer Monsoon (EASM) and East Asian Winter Monsoon (EAWM), which climatic presence were unequivocally detected in southern South China Sea (Henglai et al., 2024; Mathew et al., 2020). Collision of the Indian Plate with Asia had resulted in progressive uplift of the Tibetan Plateau during the Paleocene–Eocene, causing the Asian continent to undergo dynamic climate transitions. Significantly strengthened monsoon-inducing atmospheric circulation apparently drove accelerated amplification of the (i) Indian Summer Monsoon & Somali Jet; (ii) the Southeast Asian Monsoon, the East Asian Summer Monsoon and the East Asian Winter Monsoon; and (iii) further intensification both East Asian Monsoons; and were associated with each major orogenic pulses i.e. at (i) ca. 40–35 Ma; (ii) ca. 25–20 Ma; and (iii) ca. 15–10 Ma (Mathew et al., 2020).

Chung et al., (2014) reported that their estimated mean divergent crown age for their Clade SVLB at 8.06 mya (5.09 to 11.21 mya) coincided with the most extensive stage of karstification in Guangxi during the Miocene (Liu, 2003, 1997) and the onset and intensification of the East Asian monsoon at ca. 7.2 mya (An, 2000) to 11 mya (Zheng et al., 2004). The Sino-Vietnamese limestone Begonias evolved four times with three habits (rhizomatous; tuberous/deciduous and upright stem), with extensive karstification and it was postulated that warm (high temperature) and humid (precipitation) climate from the East Asian summer monsoon facilitated erosion of alluvium, exposing tower karsts that provided many microhabitats for rapid diversification of Begonias and other limestone karst flora (Chung et al., 2014); as well as triggered rapid diversification of both seasonally and wet adapted *Begonia* in continental Asia (Rajbhandary et al., 2011). Chung *et al*., (2014) found that rhizomatous species were most successful and proliferated in the Sino-Vietnamese limestone karst terrains, in contrast to previous suggestions that tubers and deciduous habit such as exemplified by limestone *Begonia* in Madagascar, traits associated with strong seasonal drought (Grubb, 2003; Keraudren-Aymonin, 1983). However, limestone species in Pen. Msia do not share apparent morphological trait(s) that correlates with limestone habitat, but rather facultatively possess traits suitable to grow among cracks/fissures of limestone rocks, such as attaching roots to absorb nutrients and water from thin soil layer with periodic water shortage, and able to withstand exposures to harsh terrain conditions (e.g. high temperatures, desiccation prone-), coupled with favourable reproductive traits such as small seeds to ease dispersal, non-dormant habit…*etc*. Tubers are not especially adaptive in equatorial climates, as perennial Begonias with rhizomes such as Sect. Jackia and Sect. Platycentrum proliferate as rock plants on granite- and limestone respectively. Additionally, extant limestone Begonias in Pen. Msia are not obligates, as attempts to cultivate them on other media substrates are successful provided the substrate is free draining, although watering using weak solutions of calcium carbonate significantly enhances their growth (Joanne Tan, *pers. comm*.).

## Speciation, high endemism rates of Begonias in Peninsular Malaysia, especially limestone Begonias

Many previous studies had shown that palaeogeological events and climatic fluctuations have important yet relatively different implications on speciation and diversification - influencing contemporary genetic diversity, phylogeography patterns and population structure (Li *et al*., 2015 and references therein). From the mid- to late Miocene, perhumid (wetter) climate conditions became favorable for rain forest lineages, a result of climate amenable to rain forest expansion (Crayn et al., 2015; Morley, 2012; Morley et al., 2020); however, subsequent climatic change to a Pleistocene “savannah period” might have then restricted rainforest coverage and habitats (Morley, 1998). Cave development and landscape studies in parts of Malaya suggested that higher global sea levels existed throughout much of the Tertiary, with evidence found for sea level of at least 90 m above present in the Middle Miocene, and the Pliocene (Gillieson, 2005) where evidence pointed to drier, more seasonal climates (Bird et al., 2005). These events would have provided opportunities for Begonias to conquer new or vacant niches such as on exposed limestone karsts, expanded rainforests as well as experience associated selection pressures and subsequent contractions, influencing distribution patterns of extant species- and populations. Thomas et al., (2011, 2012) had ascribed diversification/radiation of Begonias in eastern Malesia during the Pliocene and Pleistocene to formation of topographical heterogeneity and suitable microhabitats by rapid orogenesis and associated microallopatry due to pronounced sea level, climate fluctuations and associated shifts in forest distributions (Sulawesi and New Guinea).

Patterns of population structuring by locality instead of habit are reflected on the combined phylogram by populations of limestone species *B. kingiana* (widespread), *B. foxworthyi* (moderate widespread)*, B. nurii* as well as granite/sandstone species *B. sinuata* (widespread), regardless of distribution. For example, *B. foxworthyi* samples from Lata Tingkat (granite) and Hutan Simpan Pasoh (limestone), both sites located in Negeri Sembilan, grouped together and separated from a subclade of other *B. foxworthyi* (granite) samples from Lanchang, Pahang. Remainder limestone samples of *B. foxworthyi* from Kelantan and Pahang also resolved closely or found intermixed with *B. nurii* from within these locations. Samples of both *B. sinuata* and *B. kingiana* also clustered based on locality in their respective clades. A previous study on the widespread, endemic *Begonia* species, *B. maxwelliana* from Pen. Msia also found strong population differentiation (Chan et al., 2018). Similar trends were reported for African species such as *B. sutherlandii* (Hughes and Hollingsworth, 2008), *B. dregei* and *B. homonyma* (Matolweni et al., 2000), central American species *B. heracleifolia* and *B. nelumbiifolia* (Twyford et al., 2014) and *B. luzhaiensis* (Tseng et al., 2019). One main explanation was allopatric speciation where typical vicariant events due to geographical barriers such as isolation by distance, reproductive isolation (random genetic drift) and/or adaptation to prevailing (local) ecological conditions contributing to restricted gene flow, thus enforcing genetic divergence between populations; exemplified by multiple observations of microendemic Begonias (Chan et al., 2018; Hughes and Hollingsworth, 2008; Kiew, 2005; Thomas et al., 2012).

An underlying reproductive isolation factor was the limited seed dispersal capabilities of the Pen. Malaysian Begonias, given that (i) tiny seeds that are not efficiently dispersed especially for species growing on the sheltered rainforest floor (Chan et al., 2018; Hughes and Hollingsworth, 2008); (ii) seed dispersal from capsules form (e.g. *B. foxworthyi* Sect. Jackia) and *B. phoeniogramma* (Sect. Parvibegonia) is local with very narrow chance of dispersing over a wide range. Fruits of *B. foxworthyi* are dangling capsules with three equal-sized wings, and seeds fall out with small movements (“pepper pot dispersal”); whereas *B. phoeniogramma* has a “splash cup-like” fruit and seeds are released when raindrops hit the cup with great ballistic force (Kiew, 2005). It was also reported that narrow, endemic Begonias were prevalently confined to upland or montane primary rain forests (Hughes and Pullan, 2007; Thomas et al., 2012), a trend observed in majority of Begonias Sect. Platycentrum in Pen. Msia. However, none of the limestone Begonias is montane and their distributions are limited to c. 300 m asl. Paton (1957) reported that even for limestones of the same age e.g. Uralian–Permian on either side of the Main Range – on the west of the Main Range, limestone hills form a continuous platform north-south, and east-west; however on east of the Main Range, the western part of Kelantan/north Pahang limestone is similarly continuous in north–south direction but separated in an east-west direction due to thick horizons of argillaceous and pyroclastic rocks, suggesting existence of geological barriers to gene flow and peri- or parapatric speciation underlying formation of species complex *B. foxworthyi-B. nurii–B. tigrina–B. ignorata* (Kelantan-Pahang limestone). Chung et al., (2014) proposed that new species formation over time driven by strong geographic structuring could be considered an example evolution through non-adaptive radiation and genetic drift, resulting in species differing mostly in vegetative traits.

Alternatively, these species (complex) could be ascribed to hybridization, as Begonias are known to hybridize easily in cultivation. In Pen. Msia, natural hybrids occasionally occur sympatrically, such as *B. decora* x *B. venusta* (Kiew, 2003; Teo and Kiew, 1999) and *B. sibthorpioides* subsp. *machinchangensis* x *B. sinuata* (Tan et al., 2020). Since *B. nurii* and *B. foxworthyi* do overlap on some limestone karsts, there are chances of these two species hybridizing which warrants further study. Natural hybridization has an important role in evolution of *Begonia* species as well as species complexes (Peng and Chiang, 2000).

Studying a speciose genus like *Begonia* offers valuable opportunities to observe and compare different speciation patterns, as reflected on the inferred BI phylogram. Besides allopatric- and hybridization factors being plausible explanations for frequent lineage splitting, high diversity and endemism (see above) – there are contrasting species such as *B. kingiana* (limestone species, mean age = 4.62 mya; PP 0.98; HPD 2.93 – 6.43) and *B. sinuata* (sandstone & granite species, mean age = 4.12 mya; PP 1.00; HPD 2.49 – 5.88), which estimated crown age are comparatively mid-aged, have extant widespread distributions but individuals strongly grouping by locality (strong population structures implying weak geneflow) but yet taxonomically still recognizable as single species (species cohesion, low rate of lineage splitting). One interesting consideration for this observation (*B. kingiana, B. sinuata*) could be the influence of “behavioural (physiological) inertia” on niche evolution. As explained by Farallo et al., (2020) in their study of plethodontid salamanders, regulatory behaviours such as preference and (micro)habitat selection (and/or other biological factors) that match their physiological optima consequently buffered salamanders against directional selection on physiology, resulting in niche conservatism through retention of phenotypic traits and environmental preferences over long evolutionary timescales (Ackerly, 2003; Farallo et al., 2020). However, habitat selection does not explain why certain *Begonia* species should exhibit physiological conservatism, whilst others do not i.e. speciation rate varies among lineages and geographic scales. Alternatively, this inertia could be linked to genome evolution, such as eukaryote genome size variation which large proportion generally reflects content of repeated sequences, especially transposable elements (TEs). Sessegolo et al., (2016) reported signal of strong phylogenetic inertia on genome size and TE content among 26 *Drosophila* species, where phylogram results showed clustering of the smallest genomes (≤ 180 Mb) in the melanogaster (n=7; seven species) and the pseudoobscura (n=3; four species) subgroups, both possessing relatively higher speciation rates (lineage splitting). In an earlier study of genome size and species diversification, Kraaijeveld (2010) reported a general trend of large genomes either constraining speciation rate, increase extinction rate, or both. Recently, Campos-Domínguez et al., (2022) reported that *Begonia* is indeed highly variable in chromosome number and genome size, based on their research comprising of ∼20% and 4% of all *Begonia* species respectively. Genome size variation did not directly correlate with changes in chromosome number; and larger chromosome number variations were associated with some larger *Begonia* clades as well as taxonomically complex or unresolved *Begonia* sections. It was reported that Asian *Begonia* groups (C & D) have the highest number of species (as well as polyploid species), representing a 2-fold difference in chromosome number but more than 10-folds difference in genome size, suggesting variation in amount of repetitive DNA content. Dynamic changes in genome structural variation and size could potentially play an important role in *Begonia* species radiations (Campos-Dominguez et al., 2022)(and references therein). Further studies on more Asian *Begonia* species will provide deeper insights into whether smaller- or highly dynamic genomes (or both) have clear association/impact on higher speciation rates.

## Taxonomic and conservation implications for limestone Begonias in Peninsular Malaysia

Phylogenetic results showed that sectional classifications of Begonias from Pen. Msia followed mostly as proposed by Kiew (2005) and majority of these sections are in agreement with Tebbitt et al., (2006), Thomas et al., (2011, 2012), Chung et al., (2014), and Moonlight et al., (2018), with the exceptions of (i) Sect. Parvibegonia, resolved as two sister subclades similar to Moonlight et al., (2018) but not Thomas et al., (2011), Chung et al., (2014); and (ii) Sect. Diploclinium, which has species from both major clades- A and B (shown as polyphyletic by Moonlight et al., 2018). *Begonia longifolia* was formerly placed in Sect. Sphenanthera but is shown here to be nested within Sect. Platycentrum, in agreement with the conclusion of Hughes and Girmansyah (2011) that it was untenable to maintain Sect. Sphenanthera as distinct from Platycentrum.

At present, two sets of taxonomically problematic species (species complex) exist for *Begonia* of Pen. Msia based on intergradation of morphological character states observed in selected individuals, and phylogenetic results from this study based on sampling multiple individuals of the same species where resolution showed intermixing indicate that these species, as currently circumscribed, are paraphyletic, namely in clade A (i) *B. integrifolia – B. variabilis, B. integrifolia - B. elisabethae,* and *B. phoeniogramma,* which phylogenetic results corroborate only the latter as a distinct, monophyletic species; and in clade B (ii) *B. foxworthyi – B. yenyeniae, B. foxworthyi – B. tigrina, B. foxworthyi – B. nurii,* and *B. foxworthyi – B. ignorata.* Further research is ongoing to better understand the population genetics of these species (e.g. Tam et al., 2023). It is recommended that these be recognized as separate management units corresponding to their geographic lineages for conservation purposes, pending comprehensive taxonomic revision and re-circumscription.

Conservation of Begonias in Pen. Msia remains particularly concerning because they are confined to primary vegetation types and 89% are endemic (Chua et al., 2009; Kiew, 2005); and 60% are threatened of which 56% are known from fewer than five localities and 39% (21 species) from a single locality. In lowland forest, habitat loss and degradation pose the greatest threats as lowland forest habitats have been greatly reduced with an estimated 43.6% forest remaining in Peninsular Malaysia in 2018 (Tang, 2021). This was the cause of the extinction *of B. eiromischa* (Kiew, 1991, 1989). The ten Begonias that grow on limestone karsts face threats mainly from quarrying, with 75 of 445 karsts are sites of active or former quarries (Liew et al., 2016; Price, 2014) and loss of a buffer zone of trees around the karst base. Limestone forest has the highest level of extinction of any vegetation type in Pen. Msia where recent documentation includes five species from Batu Caves, Selangor (Rafidah, 2023); six species from the state of Perak (Tan et al., 2024); *Paraboea bakeri* from Gunung Sagu and Tenggek, Pahang (Rafidah and Tan, 2012). *Begonia jayaensis* is in imminent danger of extinction together with a balsam when their only known limestone habitat in Kelantan, is inundated by a hydro-electric dam currently under construction.

Only 1.6% (seven) of the 445 karst sites have legal protection because they lie within a Totally Protected Area, such as a national or state park, wildlife area or protection forest. Hence, not unexpectedly limestone forest is therefore one of the most threatened vegetation types in Pen. Malaysia (Kiew, 1997; Saw et al., 2009) and has been classified as Environmentally Sensitive Area (ESA) Rank 1 in the Malaysian National Physical Plan. ESAs require integral planning and management but currently there is no national strategy for the conservation of the biodiversity of limestone karst or for their commercial exploitation. Admittedly, limestone karsts present a specific conservation challenge because a single karst at most harbours a quarter of the limestone flora (Kiew, 2019). The best collected karst, Batu Caves in Selangor is home to 366 vascular plant species (Kiew et al., 2023) but many other hills remain poorly studied (Tan et al., 2024). As many of the endemics are range-restricted, known from a single karst (Kiew et al., 2023), this means that a network of biodiverse karsts needs to be protected to conserve maximum biodiversity, a strategy advocated by Tan et al., (2024) for the state of Perak where analysis showed protection of 15 of 95 karsts would protect 89% of the Perak’s limestone flora, 94% of its endemic species and 92% of its threatened species. There are 12 recognized national geoparks – Langkawi-Kedah, Kinabalu-Sabah (both with UNESCO Global Geopark status), and Jerai-Kedah, Kinta- & Lenggong-Perak, Labuan, Sarawak Delta-Sarawak, Strong-Kelantan, Kenyir-Terengganu, Perlis and Lipis-Pahang, with the recently established Gombak - Hulu Langat Geopark (112,955 Ha) in Selangor, which offers some level of protection to 31 geosites, including Batu Caves. However, at present there is still no national strategy nor state level mechanism for moving forward to legally protect any of these biodiverse limestone karsts or regulate their commercial exploitation.

## Conclusions

Limestone karsts form hill forests that contain unique flora (and fauna) that are highly biodiverse in comparison with the small area it occupies, constantly face threats of degradation and permanent destruction due to modern destructive practices from quarrying/mining, human encroachment and disturbances. A comparative study with thoughtfully selected species of a speciose genus like *Begonia* in Pen. Msia which shows high endemism (89%) and grows naturally on both limestone as well as non-limestone habitats has successfully provided interesting findings and significant insights into different speciation patterns and factors - such as evolution and phylogenetic relationships of Begonias, allopatric speciation (geographical barriers), reproductive isolation (traits) as well as plausible considerations for species cohesion (physiological inertia in niche evolution, genome evolution dynamics). Further research is now ongoing to resolve taxonomic status of the two sets of species complexes (paraphyletic species). More cytological studies to extend knowledge on genome size and chromosome numbers of more Asian *Begonia* species will certainly complement current research and provide holistic understanding of *Begonia* evolution. Taken together, our research contributes to increase knowledge bank of limestone flora biodiversity and evolution, which can be used to guide conservation management and practices.

## Acknowledgements

FSF would like to thank Taylor’s University for her M.Sc. scholarship. The authors are grateful to the Kepong Herbarium and Nursery (FRIM) for providing access to the Begonia living collection, notably J.P.C. Tan, P.T. Ong and the Curator of KEP. This work was partly supported by the “Flora of Peninsular Malaysia Project” (01-04-01-000 Khas) and ‘Documentation & Inventory Flora of Malaysia project’ based at Forest Research Institute Malaysia (FRIM). Special thanks also to previous Bachelor of Biotechnology (Honours) students whose Honours projects were involved in the Begonia project from the School of Biosciences, Taylor’s University Malaysia (2013 to 2020). We dedicate this paper to the late Dr. Ruth Kiew (1946 – 2025), world-renowned botanist for her steadfast dedication to Begonia research, in reference to her continuous enthusiasm over “…many more Begonia (and other) species yet to be discovered and studied…”, *souvent me souvient*.

## Author contributions

Kiew conceptualized and designed the study, planned field sampling & collected the field material, supervised the research, provided funding, and reviewed the manuscript; S.M. Tam designed the study, supported fieldwork & sampling, planned the molecular experiments, supervised the research (F. S. Fareed, L. C. Foong), provided funding, and wrote & reviewed the manuscript with input from R. Kiew; F. S. Fareed conducted molecular experiments & data analysis, and produced the figures; L. C. Foong conducted molecular experiments, reviewed and edited the manuscript, produced the tables and figures; All authors approved the final version of the manuscript.

## Supplementary material

**Supplementary file 1.** Details of 30 Begonia species (totalling 177 samples) of all seven Begonia sections from Peninsular Malaysia (sensu Moonlight et al., 2018) used in this study, including all ten limestone species, thirteen non-limestone (forest-) and seven granite species (recorded in Malaysia Biodiversity Information System (2020). **Supplementary file 2.** Forward- and reverse primer sequence information for ndhF-rpl32 and ITS DNA regions used in this study.

**Supplementary file 3.** Protocol for Polymerase Chain Reaction (PCR) reactions, performed for 40 cycles, adjusted from Thomas et al. (2011) for ndhF-rpl32 region and Chung et al. (2014) for the ITS region respectively.

**Supplementary file 4.** Maximum clade credibility chronogram estimated using BEAST. “*” represent calibration points for molecular dating. Node heights indicate mean ages and node bars indicate 95% highest posterior density (HPD).

