## Supplementary Files for "Comparative phylogenetic analyses of limestone and non-limestone Begonias in Peninsular Malaysia"

Supplementary file 1. Details of 30 Begonia species (totalling 177 samples) of all seven Begonia sections from Peninsular Malaysia (*sensu* Moonlight et al., 2018) used in this study, including all ten limestone species, thirteen non-limestone (forest-) and seven granite species (recorded in Malaysia Biodiversity Information System (2020)).

| Sections | Species | Accession Number | No. of Samples | Location of origin | Habitat | Distribution | Conservation Status | Notes |
| --- | --- | --- | --- | --- | --- | --- | --- | --- |
| *Bracteibegonia* | *B. barbellata* | FSF10 | 1 | FRIM | Non- limestone/forest | Widespread | Near Threatened | - |
| *Diploclinium* | *B. jayaensis* | FRI88520 | 4 | Gua Jaya, Kelantan | Limestone restricted | Endemic and found in only one locality | Near Threatened | - |
|  | *B. jayaensis* | FSF15 |  | Gua Jaya, Kelantan |  |  |  |  |
|  | *B. jayaensis x B. jiewhoei* | KBG2017-0918 |  | Gua Chawan, Kelantan |  |  |  |  |
|  | *B. jayaensis* | KBG2017-0917 |  | Gua Chawan, Kelantan |  |  |  |  |
| *Diploclinium* | *B. lowiana* | FRI75721 | 2 | Cameron Highlands, Pahang | Non- limestone/forest | Endemic and found in two localities | Least Concern | - |
|  | *B. lowiana* | CHRK5442 |  | Cameron Highlands, Pahang |  |  |  |  |
| *Jackia* | *B. foxworthyi* | KBG2018-0121 | 53 | Gua Sabang, Kelantan | Limestone | Endemic and moderately widespread | Near Threatened | LP1S1= Lentang Population 1 Sample 1 (granite samples) P1S1= Hutan Simpan Pasoh Population 1 Sample 1 (limestone samples) GP1S1= Lata Tingkat Popultaion 1 Sample 1 (granite samples) |
|  | *B. foxworthyi* | KBG2017-0292 |  | Gua Ikan, Kelantan | Limestone |  |  |  |
|  | *B. foxworthyi* | FRI88469 |  | Batu Baloh, Gua Musang, Kelantan | Limestone |  |  |  |
|  | *B. foxworthyi* | FRI88647 |  | Gua Hantu, Gua Musang | Limestone |  |  |  |
|  | *B. foxworthyi* | KBG2018-0223 |  | Kg. Bertam Lama, Gua Musang, Kelantan | Limestone |  |  |  |
|  | *B. foxworthyi* | FRI88776 |  | Bkt. Kemiri, Batu Baloh, Gua Musang, Kelantan | Limestone |  |  |  |
|  | *B. foxworthyi* | FRI88777 |  | Gua Berangin, Batu Baloh, Gua Musang, Kelantan | Limestone |  |  |  |
|  | **B. foxworthyi* | FRI90762 (LP1S1/S2/S3, LP2S1/S2/S3, LP3S1/S2/S3) |  | Lanchang Forest Reserve, Pahang | Granite |  |  |  |
|  | *B. foxworthyi* | FRI88981 |  | Unnamed hill, Gua Musang, Kelantan | Limestone |  |  |  |
|  | **B. foxworthyi* | FRI95859 (P1S1/S2/S3/S4, P2S1/S2/S3, P3S1/S2/S3, P4S1/S2/S3) |  | Hutan Simpan Pasoh, Negeri Sembilan | Granite |  |  |  |
|  | **B. foxworthyi* | FRI82475 (GP1S1/S2/S3) |  | Lata Tingkat, Pasoh, Negeri Sembilan | Granite |  |  |  |
|  | **B. foxworthyi* | FRI82474 (GP2S1/S2/S3) |  | Lata Tingkat, Pasoh, Negeri Sembilan | Granite |  |  |  |
|  | **B. foxworthyi* | FRI82476 (GP3S1/S2/S3) |  | Lata Tingkat, Pasoh, Negeri Sembilan | Granite |  |  |  |
|  | **B. foxworthyi* | FRI82477 (GP4S1) |  | Lata Tingkat, Pasoh, Negeri Sembilan | Granite |  |  |  |
|  | **B. foxworthyi* | FRI82478 (GP5S1/S2/S3) |  | Lata Tingkat, Pasoh, Negeri Sembilan | Granite |  |  |  |
|  | **B. foxworthyi* | FRI82479 (GP6S1/S2/S3) |  | Lata Tingkat, Pasoh, Negeri Sembilan | Granite |  |  |  |
|  | **B. foxworthyi* | FRI82480 (GP7S1/S2/S3) |  | Lata Tingkat, Pasoh, Negeri Sembilan | Granite |  |  |  |
|  | *B. foxworthyi* | FSF12 |  | Chiku 7, Gua Musang, Kelantan | Limestone |  |  |  |
|  | *B. foxworthyi* | FRI68129 |  | Lentang Forest Reserve, Bentong, Pahang | Granite |  |  |  |
|  | *B. foxworthyi* | FSF13 |  | Merapoh, Pahang | Limestone |  |  |  |
|  | *B. foxworthyi* | FSF14 |  | Chiku 8, Gua Musang | Limestone |  |  |  |
| *Jackia* | *B. ignorata B. ignorata* | FRI6008  FSF11 | 2 | Gua Senyum FRIM | Limestone restricted | Endemic and moderately widespread | Least Concern | - |
| *Jackia* | *B. nurii* | FRI88648 | 11 | Gua Musang, Kelantan | Limestone restricted | Endemic and found in two states. | Vulnerable | - |
|  | *B.cf. nurii* | FRI80821 |  | Gua Kechil, Pahang |  |  |  |  |
|  | *B. nurii* | FRI88642 |  | Gua Batu Boh, Gua Musang, Kelantan |  |  |  |  |
|  | *B. nurii* | FRI88646 |  | Gua Subong, Gua Musang, Kelantan |  |  |  |  |
|  | *B. nurii* | KBG2018-0118 |  | Gua Subong, Gua Musang, Kelantan |  |  |  |  |
|  | *B. nurii* | KBG2018-0234 |  | Gua Panjang, Gua Musang, Kelantan |  |  |  |  |
|  | *B. nurii* | FRI75462 |  | Gua Gunting, Merapoh |  |  |  |  |
|  | *B. nurii* | FSF08 |  | Chiku 7, Gua Musang |  |  |  |  |
|  | *B. nurii* | FSF09 |  | Gua Tabung, Merapoh |  |  |  |  |
|  | *B. nurii* | FRI82473 |  | Gua Gajah, Merapoh |  |  |  |  |
|  | *B. nurii* | FRI78436 |  | Gua Ulu, Kumbang, Merapoh |  |  |  |  |
| *Jackia* | *B. rajah* | FRI88682 | 2 | FRIM | Granite | Endemic and found in two states. | Vulnerable | - |
|  | *B. rajah x B. kingaiana* | KBG2018-0140 |  | FRIM | Granite |  |  |  |
| *Jackia* | *B. reginula* | FRI82472 | 1 | Ulu Senting, Negeri Sembilan | Granite and on mossy tree trunks | Endemic and found in two states. | Vulnerable | - |
| *Jackia* | *B. tigrina B. tigrina* | KBG2017-0711 FRI66588 | 2 | Gua Maka, Kelantan Gua Setir, Kelantan | Limestone restricted | Endemic and found in two localities | Critically Endangered | - |
| *Jackia* | *B. yenyeniae* | FRI47028 | 1 | Endau Rompin State Park, Segamat | Non- limestone/forest | Endemic and found in only one locality |  | - |
| *Parvibegonia* | *B. elisabethae* | KBG2012-0162 | 1 | Langkawi, Kedah | Granite or Non- limestone/forest | Found in two populations | Critically Endangered | - |
| *Parvibegonia* | *B. integrifolia B. integrifolia B. integrifolia x B. foxworthyi B. integrifolia* | FRI88945 KBG2017-0645 FSF04  KBG2009-6677 | 4 | Wang Mu, Titi Tinggi, Perlis Unknown Gua Tok Guru FRIM | Limestone and granite | Widespread | Near Threatened | - |
| *Parvibegonia* | **B. phoeniogramma* | P20181213 (P1WS1/P1WS2/P1WS3/P1WS4/S5/S6/S7;  P1NS1/S2/S3/P1NS4/S5/S6/S7/S8;  P2WS1/P2WS2/S3/S4;  P2NS1/P2NS2/P2NS3/P2NS4/P2NS5;  P3WS1) | 31 | Batu Caves, Selangor | Limestone | Endemic and found in one state | Near threatened | P1WS1= Batu Cave Population 1 With Spots (on leaves) Sample 1 (limestone samples with spots on leaves) P1NS1= Batu Cave Population 1 No Spots (on leaves) Sample 1 (limestone samples without spots on leaves) |
|  | **B. phoeniogramma* | FRI90761 (GP1S1/ GP1S2; GP2S1/GP2S2/GP2S3) | Hulu Langat, Selangor | Gabai Waterfall, | Granite |  |  |  |
|  | *B. phoeniogramma* | FRI78123 |  | Batu Caves, Selangor | Limestone |  |  |  |
| *Parvibegonia* | *B. sibthorpioides* | FRI75877 | 4 | Gua Jerai, Kedah | Sandstone | Extremely rare | Critically Endangered | - |
|  | *B. sibthorpioides* | FRI80843 |  | Gunung Machinchang, Langkawi | Sandstone |  |  |  |
|  | *B. sibthorpioides* | FRI80843-1 |  | Gunung Machinchang, Langkawi | Sandstone |  |  |  |
|  | *B. sibthorpioides x B. sinuata* | _ |  | Gunung Machinchang, Langkawi | Sandstone |  |  |  |
| *Parvibegonia* | *B. sinuata* | FSF01 | 17 | Penang Hill | Granite | Widespread | Least Concern | P1S1= Sekayu Population 1 Sample 1 (granite samples) |
|  | *B. sinuata* | FSF02 |  | Gunung Machinchang, Langkawi | Sandstone |  |  |  |
|  | *B. sinuata* | FSF03 |  | Gunung Machinchang, Langkawi | Sandstone |  |  |  |
|  | **B. sinuata* | FRI82481 (P1S1/P1S2/P1S3/P1S4) |  | Sekayu Recreational, Forest, Kuala Berang, Terengganu | Garnite |  |  |  |
|  | **B. sinuata* | FRI82482 (P2S1/P2S2/P2S3) |  | Sekayu Recreational, Forest, Kuala Berang, Terengganu | Granite |  |  |  |
|  | **B. sinuata* | FRI82483 (P3S1/P3S2/P3S3) |  | Sekayu Recreational, Forest, Kuala Berang, Terengganu | Granite |  |  |  |
|  | **B. sinuata* | FRI82484 (P4S1/S2/S3) |  | Sekayu Recreational, Forest, Kuala Berang, Terengganu | Granite |  |  |  |
|  | *B. sinuata* | FRI75719 |  | Sg. Pisang, Selangor | Granite |  |  |  |
| *Parvibegonia* | *B. thaipingensis* | FRI80835 | 2 | Hulu Gombak, Selangor | Non- limestone/forest | Endemic and moderately widespread | Near Threatened | - |
| *Parvibegonia* | *B. variabilis A & B* | KBG2018-0147 | 5 | Wang Mu F.R., Perlis | Limestone | Moderately widespread | Least Concern | A & C= Big spots on leaves B & D= Small spots on leaves |
|  | *B. variabilis C & D* | FRI88061 |  | Lata Puteh, Perak | Granite |  |  |  |
|  | *B. variabilis* | KBG2018-2035 |  | Gua Panjang, Gua Musang | Limestone |  |  |  |
| *Petermannia* | *B. holttumii* | FRI55986 | 1 | Cameron Highlands, Pahang | Non- limestone/forest | Widespread | Near Threatened | - |
| *Petermannia* | *B. jiewhoei B. jiewhoei B. jiewhoei* | FRI78099 FRI70623 FRI80999 | 3 | Gua Musang, Kelantan Gua Musang, Kelantan Gua Musang, Kelantan | Limestone restricted | Endemic and found in one locality. | Critically Endangered | - |
| *Petermannia* | **B. wrayi* | FSF 05 | 5 | Lata Tingkat, Negeri Sembilan | Non- limestone/forest | Widespread | Near Threatened | - |
|  | **B. wrayi* | FSF 06 |  | Lata Tingkat, Negeri Sembilan |  |  |  |  |
|  | *B. wrayi* | FRI80702 |  | Temenggor Dam, Perak |  |  |  |  |
|  | *B. wrayi* | FRI75717 |  | Ulu Bertam, Gombak |  |  |  |  |
|  | *B. wrayi* | FRI63017 |  | G. Inas Forest Reserve, Baling, Kedah |  |  |  |  |
| *Platycentrum* | *B. abdullahpieei* | FRI64721 | 1 | Sg. Lata Puteh, Bintang Hijau FR, Larut | Granite | Endemic and found in one locality. | Critically Endangered | - |
| *Platycentrum* | *B. decora* | FRI75723 | 2 | Cameron Highlands, Pahang | Non- limestone/forest | Widespread | Near Threatened | - |
|  | *B. venusta x B. decora* | FRI75724 |  | Cameron Highlands, Pahang |  |  |  |  |
| *Platycentrum* | *B. herveyana* | FRI49292 | 1 | Bukit Senggeh, Melaka | Granite | Endemic and found in two states, | Critically Endangered | - |
| *Platycentrum* | *B. maxwelliana* | FRI70306 | 1 | Bukit Larut, Perak | Granite or Non- limestone/forest | Endemic and widespread. | Near Threatened | - |
| *Platycentrum* | *B. pavonina* | FRI75725 | 1 | Cameron Highlands, Pahang | Non- limestone/forest | Endemic and found only in one state | Vulnerable | - |
| *Platycentrum* | *B. rheifolia* | FSF07 | 1 | Rimba Ilmu, Selangor | Granite | Endemic and moderately widespread | Least Concern | - |
| *Platycentrum* | *B. rhoephila B. rhoephila* | FRI75319 FRI75718 | 2 | Pulau Tioman, Pahang Sg. Pisang, Selangor | Granite | Endemic and found in only one state. | Endangered | - |
| *Platycentrum* | *B. venusta* | FRI75722 | 1 | Cameron Highlands, Pahang | Non- limestone/forest | Endemic and widespread | Least Concern | - |
| *Ridleyella* | *B. kingiana* | KBG2018-0121 | 14 | Wang Mu F.R., Perlis | Limestone restricted | Widespread | Endangered | - |
|  | *B. kingiana* | FSF16 |  | Langkawi, Kelantan |  |  |  |  |
|  | *B. kingiana* | KBG2017-0212 |  | Kuala Bertis, Kelantan |  |  |  |  |
|  | *B. kingiana* | FRI88643 |  | Gua Batu Boh, Kelantan |  |  |  |  |
|  | *B. kingiana* | FRI88687 |  | Gua Musang, Kelantan |  |  |  |  |
|  | *B. kingiana* | KBG2018-0222 |  | Kg. Bertam Lama,  Gua Musang, Kelantan |  |  |  |  |
|  | **B. kingiana 1,2 & 3* | K20181213 |  | Batu Caves, Selangor |  |  |  |  |
|  | *B. kingiana* | FRI88984 |  | Unnamed Hill, Gua Musang, Kelantan |  |  |  |  |
|  | *B. kingiana* | FRI88989 |  | Gua Jinjang, Pelamin, Merapoh, Pahang |  |  |  |  |
|  | *B. kingiana* | FRI78124 |  | Gunung Kanthan, Perak |  |  |  |  |
|  | *B. kingiana* | KBG2009-0310 |  | Langkawi, Kedah |  |  |  |  |
|  | *B. kingiana* | FSF17 |  | Merapoh, Pahang |  |  |  |  |
| *Sphenanthera* | *B. longifolia B. longifolia* | FRI75726 FRI75728 | 2 | Cameron Highlands, Pahang | Non- limestone/forest | Widespread | Near Threatened | - |

* Samples collected from field

**Supplementary file 2.** Forward- and reverse primer sequence information for ndhF-rpl32 and ITS DNA regions used in this study.

| DNA region | Primers | Annealing Temperature | Sequences (5 → 3’) | Reference |
| --- | --- | --- | --- | --- |
| ndhF-rpl32 | ndhFBeg-F (Forward) | 50 ˚C | TGGATGTGAAAGACATATTTTG CT | Thomas et al., (2011) |
|  | trnLBeg-R (Reverse) |  | TTTGAAAAGGGTCAGTTAATAA CAA | Thomas et al., (2011) |
| ITS | 5P* (Forward) | 55 ˚C | GGAAGGAGAAGTCGTAACAAGG | Clementet al., (2004) |
|  | 26S1Rev* (Reverse) |  | CGCCTGACCTGGGGTCG | Clementet al., (2004) |

*Clement, W.L. *et al.* (2004) “Phylogenetic position and biogeography of Hillebrandia sandwicensis (Begoniaceae): a rare Hawaiian relict,” *American Journal of Botany*, 91(6), pp. 905–917.

*Thomas, D.C. *et al.* (2011) “A non-coding plastid DNA phylogeny of Asian *Begonia* (Begoniaceae): Evidence for morphological homoplasy and sectional polyphyly,” *Molecular Phylogenetics and Evolution*, 60(3), pp. 428–444. https://doi.org/10.1016/j.ympev.2011.05.006.

**Supplementary file 3.** Protocol for Polymerase Chain Reaction (PCR) reactions, performed for 40 cycles, adjusted from Thomas et al. (2011) for ndhF-rpl32 region and Chung et al. (2014) for the ITS region respectively.

| PCR Protocol | Primers | Annealing Temperature |
| --- | --- | --- |
| Initial denaturation | 80 ˚C 🡪 5 min | 94 ˚C 🡪 5 min |
| Denaturation | 95 ˚C 🡪 1 min | 94 ˚C 🡪 5 min |
| Primer annealing | 50 ˚C 🡪 1 min | 55 ˚C 🡪 5 min |
| Extension | 65 ˚C 🡪 2 min | 72 ˚C 🡪 5 min |
| Final extension | 65 ˚C 🡪 5 min | 72 ˚C 🡪 5 min |

*Thomas, D.C. *et al.* (2011) “A non-coding plastid DNA phylogeny of Asian *Begonia* (Begoniaceae): Evidence for morphological homoplasy and sectional polyphyly,” *Molecular Phylogenetics and Evolution*, 60(3), pp. 428–444. https://doi.org/10.1016/j.ympev.2011.05.006.

* Chung, K.-F., Leong, W.-C., Rubite, R.R., Repin, R., Kiew, R., Liu, Y., Peng, C.-I., 2014. Phylogenetic analyses of Begonia sect. Coelocentrum and allied limestone species of China shed light on the evolution of Sino-Vietnamese karst flora. *Boanical. Studies,* 55, 1. https://doi.org/10.1186/1999-3110-55-1

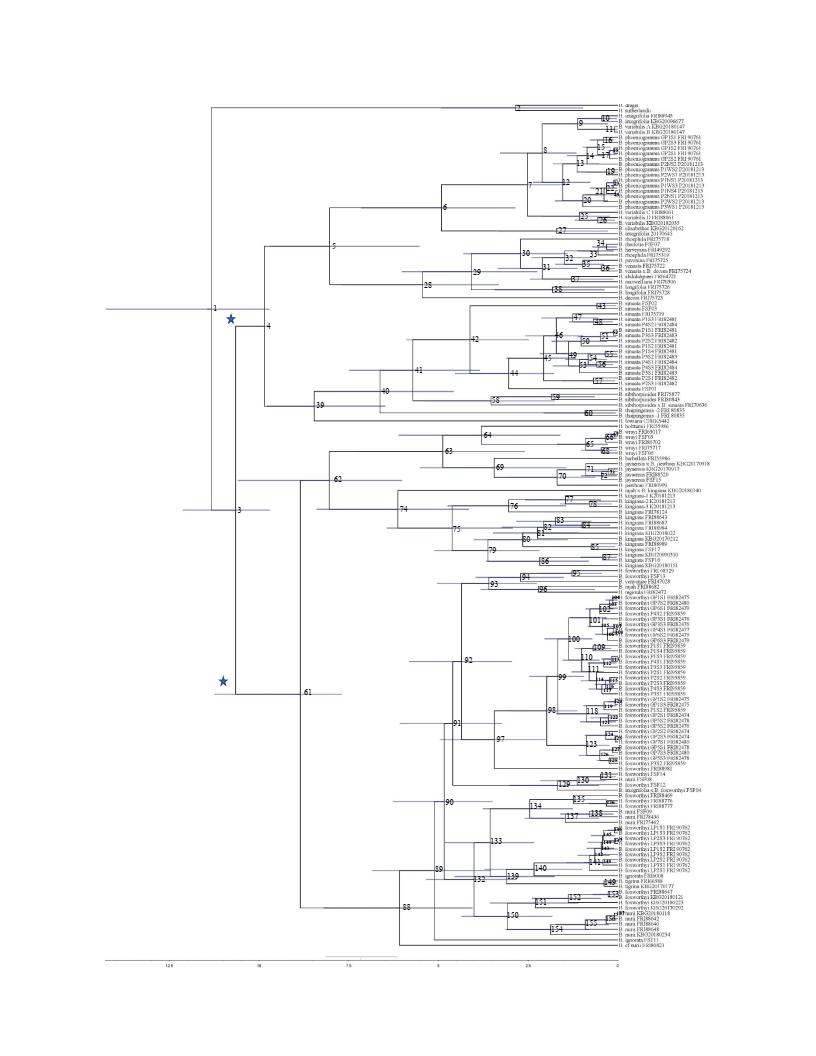

**Supplementary file 4.** Maximum clade credibility chronogram estimated using BEAST. “*” represent calibration points for molecular dating. Node heights indicate mean ages and node bars indicate 95% highest posterior density (HDP).
