## Supplementary Tables for "Comparative phylogenetic analyses of limestone and non-limestone Begonias in Peninsular Malaysia"

Table 1: Sequence information for chloroplast ndhF-rpl32 (161 samples), nuclear ITS (156 samples) and the two combined DNA regions for Begonia samples used in this study. Characters were unordered, had equal weight and gaps were treated as missing data

| Partition | Length variation (bp) | No. of aligned position (bp) | Variable characters (%) | Parsimony informative character (%) | No. of characters excluded (%) |
| --- | --- | --- | --- | --- | --- |
| ndhF-rpl32 | 800-990 | 1037 | 609 (50) | 383 (31.7) | 168 (13.9) |
| ITS | 700-800 | 833 | 632 (75.8) | 498(59.7) | 188 (18.4) |
| Combined | 1400-2000 | 1940 | 1356 (69.8) | 1086 (55.9) | 221 (10.2) |
| ndhF-rpl32+GenBank | 950-1050 | 1226 | 838 (68.3) | 636 (51.8) | 175 (12.4) |
| ITS+GenBank | 800-920 | 915 | 842 (92.0) | 684 (75.7) | 246 (26.8) |

Table 2: Maximum clade credibility chronogram and summary of divergence time estimates, and clade supports (PP) generated by the BEAST software for Peninsular Malaysian *Begonia* species

| Source | Sampling | Markers Used | Clades and Sections |
| --- | --- | --- | --- |
| Moonlight et al., (2018) | 574 species- Asian, 51 African and 283 American | ndhF-rpl32, ndhA, rpl32-trnL | Asian Begonia resolved in two clades- C and D **Clade C:** PAR, DIP, PLA, SPH **Clade D:** COE, PET, RIDL, JAC, BRAC, SYM, BARY |
| Chung et al., (2014) | 94 species | rpl16 and ITS | Asian Begonia resolved in two clades- C and D **Clade C:** PAR, DIP, PLA, SPH, REI, BARY **Clade D:** COE, PET, RIDL |
| Tebbitt et al., (2006) | 46 Asian Begonia | ITS and 5.8S | Asian Begonia resolved in two clades- *Petermannia-Coelocentrum* and *Platycentrum*-*Sphenanthera* ***Petermannia*-*Coelocentrum*:** PLA, SPH, DIP, PET, PAR, BARY ***Platycentrum-Sphenanthera:*** DIP, PET, RID |
| Thomas et al., (2012) | 112 species | ndhF-rpl32, ndhA, rpl32-trnL | Asian Begonia resolved in two clades- C and D **Clade C:** REI, PLA, PAR, DIP, SPH **Clade D:** COE, REI, BRAC, PET |
| Thomas et al., (2011) | 84 species | ndhF-rpl32, ndhA, rpl32-trnL | Asian Begonia resolved in two clades- C and D **Clade C:** REI, PAR, DIP, PLA-SPH **Clade D:** COE, REI, PET, DIP, BRAC |

BARY= *Baryandra,* BRAC= *Bracteibegonia*, COE= *Coelocentrum,* DIP= *Diploclinium,* JAC= *Jackia,* PAR=*Parvibegonia,* PET= *Petermannia,* PLA= *Platycentrum,* REI= *Reichenheimia,* RIDL= *Ridleyella,* SYM= *Symbegonia*
